# Modeling genome-wide evolution of catalytic turnover rates: Strong epistasis shaped modern enzyme kinetics

**DOI:** 10.1101/318972

**Authors:** David Heckmann, Daniel C. Zielinski, Bernhard O. Palsson

## Abstract

Systems biology describes cellular phenotypes as properties that emerge from the complex interactions of individual system components. Little is known about how these interactions have affected the evolution of metabolic enzymes. To address this question, we combine genome-scale metabolic modelling with population genetics models to simulate the evolution of enzyme turnover numbers (*k_cat_*s) from a theoretical ancestor with inefficient enzymes. This systems view of biochemical evolution reveals strong epistatic interactions between metabolic genes that shape evolutionary trajectories and influence the magnitude of evolved *k_cat_*s. A small number of biophysically constrained enzymes suffice to induce diminishing returns epistasis that prevents enzymes from developing higher *k_cat_*s in all reactions and keeps the organism far from the potential fitness optimum. In addition, multifunctional enzymes cause synergistic epistasis that slows down adaptation. The resulting fitness landscape is smooth and causes *k_cat_* evolution to be convergent. Predicted *k_cat_* parameters show a significant correlation with experimental data on *in vitro* and *in vivo* turnover rates, validating our modelling approach. Our analysis thus suggests that enzyme evolution can be predicted on a genome scale and reveals the mechanisms by which evolutionary forces shape modern *k_cat_*s and the whole of cell metabolism.

## Introduction

The biological systems we observe today are the results of evolutionary trajectories that were shaped by their underlying genotype-to-fitness map, termed the fitness landscape. The components of the system constantly change to increase fitness in the current environment. It is thus tempting to assume that, given the right environment, biological systems can be described as the state that results in the highest fitness possible under all biophysical constraints. While such optimality assumptions were successfully applied to understand a variety of systems properties like bacterial growth rates^1,2^, gene expression patterns^3-5^, and metabolic fluxes^6,7^, they are expected to prove futile when the underlying fitness landscape is rugged and exhibits local optima^8,9^, or when the natural selection cannot overcome genetic drift to establish potential fitness gains^10-12^. The topography of the fitness landscape is determined by epistasis, i.e., the extent to which the fitness effect of a mutation depends on the genetic background. Understanding epistasis is thus crucial for understanding evolutionary dynamics and constraints, and systems models can serve as a key tool to understand these interactions^9,13,14^.

It was suggested that the kinetic turnover rates (*k_cat_*s) of metabolic enzymes constitute an example of a system state that is distant from a potential optimum, as the efficiency of most enzymes remains far from its theoretical maximum^15,16^. Enzyme turnover numbers span over six orders of magnitude and are essential for understanding biological processes on a quantitative level, as they quantitatively describe the proteomic demands of reaction flux, growth, and thus fitness^2, 17-22^. In contrast to this high variability and functional importance, experimental data on *k_cat_* is scarce and exhibits high noise^15^. An improved understanding of the evolutionary and biophysical forces that shape the distribution of kinetic parameters on a systems scale would thus constitute an important step towards quantitative understanding of cellular metabolism. A meta-analysis of databases of *k_cat_*s showed two major patterns^15^. On the one hand, *k_cat_*s in primary metabolism are consistently higher than those in pathways of secondary metabolism, a finding that can be interpreted as the result of differential selection pressure on the respective genes. On the other hand, the underlying biochemical mechanism has a measurable effect on *k_cat_*, suggesting that an interplay between biophysical and evolutionary constraints determines metabolic *k_cat_*s. How these factors have acted mechanistically to result in the diverse kinetic turnover rates we observe today is unknown.

The study of evolution is often limited to retrospective phylogenetic analysis of genome sequences. Nevertheless, when the selective advantage conferred by a metabolic system can be identified, quantitative models can be used to predict fitness correlates and evolution. In the past, systems models of metabolism have been used successfully to describe a variety of evolutionary phenomena like the dynamics of genome reduction^23^, properties of ancient metabolism^24^, the global optimum of metabolic adaptation^1^, and the trajectories of photosynthesis evolution^25^. In this study we aim to understand the evolutionary mechanisms that underlie *k_cat_* evolution and its apparent failure to reach optimality. As *k_cat_*s provide a quantitative link between proteome costs and metabolic flux, metabolic models can be used to predict how *k_cat_*s affect growth as a proxy for fitness. To this end, we combine genome scale modelling of metabolism with population genetics models to simulate how modern *k_cat_*s evolved from slow ancestors in a network context.

## Results

### A model for simulating systems-wide *k_cat_* evolution

As *k_cat_*s affect fitness by controlling the proteomic cost of enzyme reactions^2,17,18,26^, we hypothesize that genome-scale models of cell growth can be used to retrace *k_cat_* evolution in a network context.

The core structure of the metabolic network is conserved across the tree of life^27,28^, and thus modern metabolic networks can be expected to contain information about the network context in which enzymes evolved. Because of the quality of its metabolic reconstruction and the relatively high coverage of kinetic data, we choose the metabolic network of *E. coli* K-12 MG1655 as an ideal candidate to study *k_cat_* evolution.

To predict *k_cat_*-dependent growth as a proxy for fitness, we use the MOMENT algorithm^4^ and a genome-scale reconstruction of *E. coli* metabolism^29^. The MOMENT algorithm optimizes growth under a constraint on the total metabolic proteome a cell can sustain. As changes in gene expression can be achieved by the gene regulatory network of the cell or through mutations in a genetic target that is much larger than that for kinetic parameter evolution, we model gene expression as growth-optimal.

Modern enzymes exhibit relatively high substrate specificity, but are assumed to have evolved from slow multifunctional ancestors^30-32^. We aim to model adaptation of kinetic turnover rates after specificity increased, but where turnover rates were still low. We thus assign turnover rates of 10^−3^ s^−1^, similar to the slowest enzymes observed today^15^. Starting from these ancestral slow enzymes, mutations are drawn randomly as multiplicative changes in *k_cat_*s of a random reaction, where the majority are assumed to be decreasing *k_cat_* (decreasing:increasing = 100:1, see Figure 1 and Methods). Whether a novel mutation achieves fixation is then calculated for the estimated effective population size of *E. coli* (*N_e_* = 2.5e7,^33,34^), and *k_cat_* evolution is simulated with a Markov Chain Monte Carlo approach (MCMC, Figure 1). The model thus uses a strong-selection-weak-mutation regime^35^.

**Figure 1:**
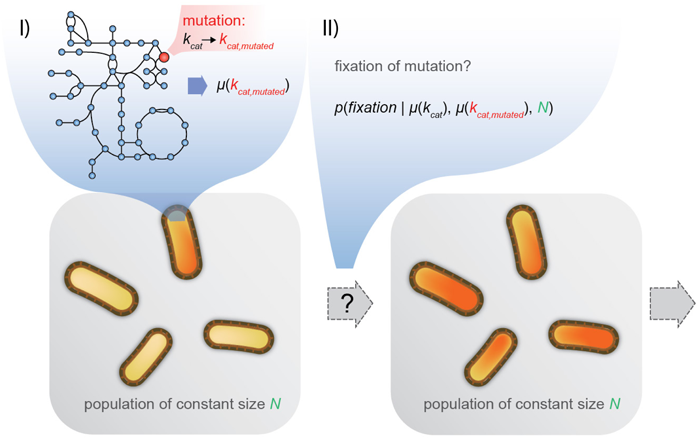
A single iteration of the MCMC algorithm used for simulating genome-scale *k_cat_* evolution: (I) A mutation in the *k_cat_* of a random reaction of a single cell in the population is introduced. The original growth rate *μ(k_cat_)* and the novel growth rate *μ(k_cat,mutated_)* are predicted by solving the respective MOMENT problems (see Methods). (II) The probability of fixation for the novel mutation is calculated with a population genetics model based on *μ(k_cat_)*, *μ(k_cat,mutated_)*, and the population size *N*. Fixation of the novel change in *k_cat_* is then decided based on this probability. If fixation fails, the mutation is discarded. A typical simulation run includes around 10^8^ of the described iterations.

As biological catalysts are limited to natural amino acids to stabilize transition states, it is expected that many reactions will have a distinct biophysical upper limit to the turnover rate that is lower than the theoretical limit resulting from diffusion rate of collisions. As certain reaction mechanisms were shown to consistently exhibit high *k_cat_*s^15^, we use the enzyme commission (EC) number to decide on a candidate set of 569 biophysically unconstrained reactions (see Methods). The remaining 1087 enzymes were considered biophysically constrained and were fixed to the median of *in vitro k_cat_* measurements (13.7 s^−1^). In the context of evolutionary predictions, the number of enzymes in the constrained and unconstrained set are more meaningfully compared in terms of reactions that are contributing to growth. Flux variability analysis^36^ for aerobic growth on glucose reveals that 278 growth-relevant reactions (see Methods) are unconstrained, while 183 carry a biophysical constraint; the majority of *in silico* growth-relevant reactions is thus evolving without upper limit.

### Evolutionary trajectories exhibit jumps, convergence and diminishing returns

When simulating *k_cat_* evolution with the MCMC algorithm, we can trace the dynamics of adaptation through evolutionary trajectories of growth rates (Figure 2A). As a starting point, we choose an aerobic glucose environment. Ancestral slow enzymes cause initial growth rates to be low, but fixation of mutations that increase selected *k_cat_*s leads to an irregular increase in growth rates that eventually saturates towards a growth rate close to 0.5 h^−1^. This behavior is reproducible across replicates, and final growth rates are convergent across these independent evolutionary trajectories. Even though the majority of growth-contributing reactions—as determined by flux variability analysis^36^—were not assigned biophysical constraints on the evolution of higher *k_cat_*s, growth rates are unable to surpass 0.5 h^−1,^ even when simulations are continued further than shown in Figure 2 (Supplemental Figure 2, Supplemental Figure 4). This effect is the result of diminishing returns epistasis (DRE) acting between the evolving genes: the same mutation will result in a smaller fitness gain when the genetic background already enables a high growth rate (Figure 2C). Due to this effect, even large improvements in *k_cat_*s of high-flux pathways can only confer a fitness benefit that approaches that of a neutral mutation and thus become subject to drift rather than selection^10,11^ (Figure 2B). We confirm this idea by using a greedy search that iteratively fixes the most beneficial mutations that double kcat: the maximum achievable fitness gain will reach the neutral barrier (where *s* is smaller ^~^1/*N_e_* ^10,11^) without achieving a growth rate greater than 0.5 h^−1^ (Supplemental Figure 4). The underlying mechanism for the observed DRE is the dispersion of biophysical constraints through the shared metabolic proteome (Supplemental Information 2); as genome-wide adaptation progresses, improvements of already high *k_cat_*s free up little protein that can be invested in limited reactions. This effect is independent of whether multiplicative or additive mutations are used (Supplemental Information 2). We simulated a maximum growth rate that ignores evolutionary constraints by setting the *k_cat_* of all unconstrained reactions to a value similar to the fastest known enzymes of 1e5 s^−1^. We find a theoretically achievable growth rate of 1.58 h^−1^, more than three times the rate of the evolved result. This result indicates the strong effect that DRE has in constraining *k_cat_* evolution.

**Figure 2:**
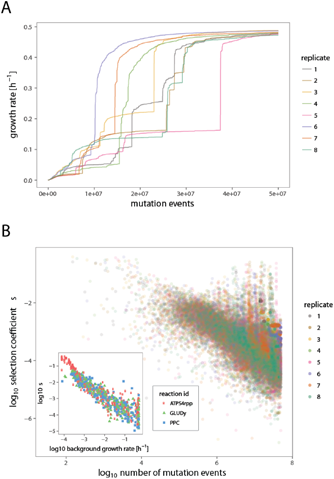
Evolutionary trajectories and diminishing returns epistasis: (A) The growth rate of the population against the number of simulated mutations. The replicates showed 4880 fixation events on average. (B) The selection coefficient *s* (defined as the change in growth rate relative to the novel growth rate) against the cumulative number of simulated mutational events for all fixed mutations. The inset shows *s* against the background growth rate in which a mutation occurred for the three reactions that had the most changes in *k_cat_* fixed. These reactions are: ATPS4rpp: ATP synthase, GLUDy: Glutamate dehydrogenase (NADP), and PPC: Phosphoenolpyruvate carboxylase.

While *in vitro* data and biochemical reaction mechanisms defined our set of biophysically constrained reactions, the true identity of this set is unknown. We thus conducted a sensitivity analysis for the identity and size of this set. The identity of the evolving set affects the final growth rate, but not the qualitative dynamics of adaptation or the occurrence of DRE (Supplemental Figure 5). The speed of adaptation decreases with the size of the evolving set, as more reactions are required to acquire mutations to reach higher growth rates. An additional source of uncertainty comes from the nature of the distribution of mutational effects, which is unknown. We varied the mean of the distribution of mutational effects, but again found no effect on the qualitative dynamics of adaptation or the occurrence of DRE, but a small quantitative effect on the final growth rate (Supplemental Figure 6).

### Multifunctional enzymes cause evolutionary jump dynamics

In order to understand the irregular increase in growth rate observed in adaptive trajectories (Figure 2A), we summarized genes for which mutation coincided with unusually high fitness gains. We found a small set of genes that was repeatedly responsible for large jumps in fitness (Supplemental Table 1). Investigation of metabolic network model and gene-protein-reaction context of these genes revealed that all of them are multifunctional enzymes that catalyze multiple reactions in the same linear pathways. These genes are involved in histidine biosynthesis (*histb*), purine biosynthesis (*purH*), and cell wall biosynthesis (*glmU*), and fatty acid biosynthesis (*fabG*). The irregular behavior in adaptive trajectories thus has a mechanistic reason that lies in the structure of the underlying network: protein cost of the linear pathway cannot be mitigated by increasing an individual *k_cat_* of a single active site, resulting in a fitness landscape that shows synergistic epistasis (Figure 1, Figure 3). The pathway can then become a bottleneck for the adaptation process, where fixation of a specific neutral mutation in a multifunctional enzyme is required for further fitness gains (Figure 3B).

**Figure 3:**
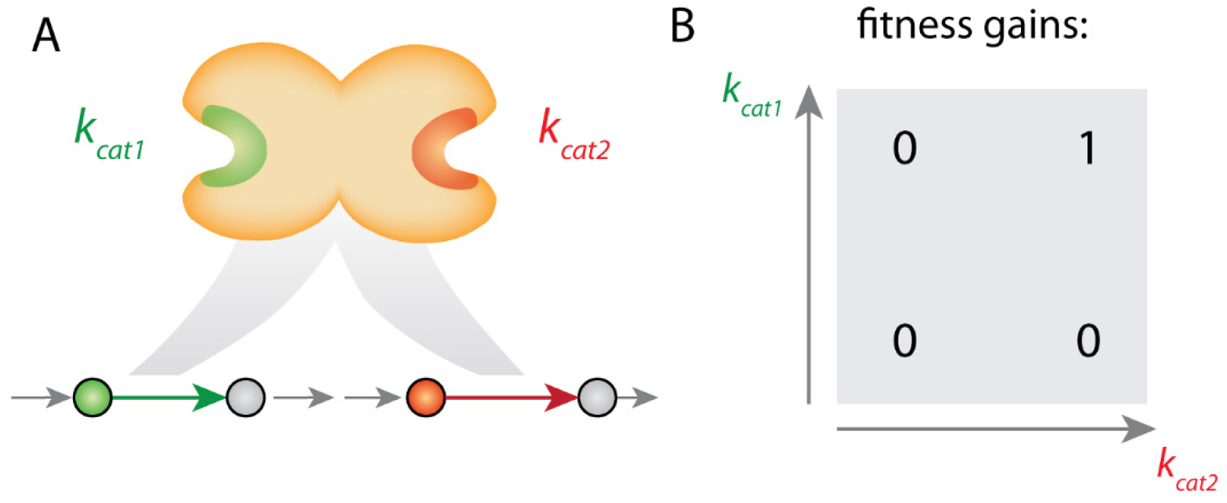
Multifunctional enzymes cause synergistic epistasis in *k_cat_* evolution: (A) A multifunctional enzyme with two distinct active sites catalyzes two reactions in the same linear fitness-relevant pathway. (B) Mutations that increase either *k_cat_* individually cannot be used to reduce protein cost of the pathway and thus exhibit synergistic epistasis.

### Most, but not all, reactions show repeatable evolution

The high level of convergence that is exhibited in the adapted growth rates (Figure 2A) is reflected in the turnover rates of the evolved populations: vectors of adapted *k_cat_*s show a high correlation across replicates (all Pearson’s *R^2^*>=0.9, Supplemental Figure 1). Clustering of the most divergent reactions reveals that the remaining differences in evolved *k_cat_*s cannot be exclusively attributed to the stochasticity of the adaptation process: redundant metabolic routes in central carbon metabolism and redox metabolism cause *k_cat_* evolution to be divergent (Supplemental Figure 1B). Nevertheless, *k_cat_* evolution is highly convergent and repeatable, indicating that similar patterns in turnover rates across species could be the result of independent evolutionary trajectories.

### The evolved vector of *k_cat_*s agrees with *in vivo* and *in vitro* data and considering diverse environments improves the agreement

How well do our simulated end points of *k_cat_* evolution agree with experimental data on modern *k_cat_*s? In order to answer this question, we simulate *k_cat_* evolution in randomly changing model environments to model a more realistic environmental diversity. We randomly chose a set of environmental carbon, nitrogen, and sulfur components, as well as random availability of oxygen (see Methods) and compared prediction performance of this diverse environment simulation with the simulations under constant aerobic glucose conditions.

*In vitro* measurements of *k_cat_* were previously mined from the BRENDA database and filtered for natural substrates^15^. We compared the simulated end points for both constant and diverse environments to this data set while focusing on reactions without data-driven biophysical constraints to avoid circular conclusions. We found that the predictions agree in magnitude (Figure 4A, Supplemental Figure 7A) and show a significant correlation (Pearson’s *R*=0.37, *p*<6e-4 for diverse environments. *R*=0.25, *p*<0.02 for aerobic growth on glucose. See Methods) with the *in vitro* data (Figure 4B, Supplemental Figure 7B). Simulation of evolution in diverse environments thus results in a better agreement with *in vitro* data. In addition to *in vitro* measurements, estimates of *in vivo* maximal turnover rates (*k_app,max_*) became recently available based on the combination of proteomics data and flux predictions across multiple conditions^37^. The predicted *k_cat_*s from both diverse and constant evolutionary environments agree with this *in vivo* data in magnitude (Figure 4C, Supplemental Figure 7C) and show a highly significant correlation (*R*=0.67, *p*<5e-29, for diverse environments. *R*=0.57, *p*<2.4e-19 for aerobic growth on glucose). Like in the case of *in vitro* measurements, a model of diverse environments explains *in vivo* data better than constant environments.

**Figure 4:**
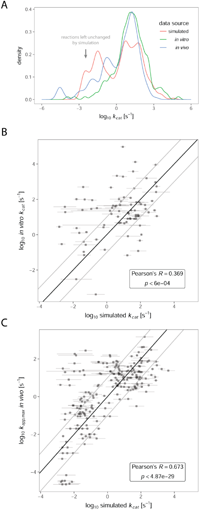
Comparison between *k_cat_* predictions for evolution in diverse environments and experimental data. (A) Distribution of turnover rates in *in vitro* (n=188)^15^, *in vivo* (n=210)^37^, and simulated data (n=276). Simulated data is only shown for non-constrained reactions that contribute to growth. The arrow indicates reactions that were essentially left unchanged by the simulation, indicating that they were not used in most environments. (B) Comparison between experimental *in vitro* data and simulated data for reactions contributing to growth (n=83). (C) Comparison between experimental *in vivo* data (*k_app,max_*) and simulated data for reactions contributing to growth (n=210). The outliers in the upper left suggest that these reactions are rarely used in the environmental conditions that we model. Horizontal error bars in (B) and (C) show the standard deviation across three simulated replicates.

## Discussion

The turnover rates of enzymes in central metabolism are significantly higher than those of pathways in secondary metabolism^15^, even though phylogenetic evidence suggest that the core of the metabolic network is conserved across the tree of life^27,28^ and extensive enzyme optimization should thus have had sufficient time to occur. In order to understand the mechanistic reason for this observation, we developed an *in silico* model that predicts the dynamics and long-term end point of *k_cat_* evolution, and validated these predictions with experimental data.

### Diminishing returns epistasis prevents *k_cat_* optimization

It has been suggested that the suboptimal turnover rate of many enzymes is the result of an increasing difficulty to achieve *k_cat_* improvements that occurs in all metabolic genes^15^. We show that even without such intragenic constraints, a small number of biophysically constrained reactions are sufficient to cause diminishing returns epistasis in otherwise unconstrained reactions (Figure 2, Supplemental Figure 5, Supplemental Information 2). As the fitness gain of improvements in *k_cat_*s (i.e., their selection coefficient *s*) decreases, it approaches the neutral boundary that lies around 1/*N_e_*^10,11,38^, and mutations that yield large improvements in *k_cat_* are rendered effectively neutral.

Metabolic control theory^39^ has been used in the past to postulate the occurrence of diminishing fitness returns when the activity of a single enzyme changes, e.g., explaining the genetic dominance of metabolic genes^40^ and the frequency of neutral mutations^38^. Such diminishing returns that result from the adjustment of single enzyme activities are not the source of the diminishing returns we report for the metabolic macro-evolution at genome-scale: When biophysical constraints are removed from our model, the growth rate expands without bounds (Supplemental Figure 3), showing that these constraints cause diminishing fitness gains in our model.

A mechanism of intergenic diminishing returns epistasis between beneficial mutations that alleviate the expression costs of metabolic enzymes was found in experimental evolution^41^: the expression cost of a heterologous pathway was reduced by reducing over-expression, a process conceptually similar to the reduction of protein costs through the increase in kinetic parameters. While the adjustment of expression levels is a mechanism commonly found in experimental evolution, kinetic parameter evolution is a smaller mutational target and thus more difficult to study in such a framework.

### Convergent evolution on a smooth fitness landscape

Structural genomics studies have found convergent evolution of function to be a common pattern in enzyme evolution^42^. Our model shows that kinetic parameter evolution is likely to similarly exhibit convergent behavior. The evolutionary end points show a high correlation of *k_cat_*s across replicates— even though some reactions diverge—(Supplemental Figure 1), and final growth rates are very similar (Figure 2). This suggests a smooth single-peaked phenotypic fitness landscape, where the low level of divergence indicates a plateau of comparable fitness that is reached in a repeatable and convergent manner. Remarkably, this high level of convergence is even found when environments differ during the adaptation process (Figure 4). Even though diminishing returns epistasis arises for the growth rate effect of mutations, epistatic effects of mutations in the same gene are not modelled explicitly. Thus, even though structural models argue against this^43^, intragenic sign epistasis—where the sign of a mutation’s effect depends on the genetic background—could cause a more rugged landscape.

Although the model suggests a remarkably smooth fitness landscape, multifunctional enzymes cause “neutral plateaus” that slow adaptation by requiring a neutral mutation to occur before *k_cat_* improvements can yield fitness gains (Figure 3). Most of these cases are caused by multifunctional enzymes that possess two distinct active sites and that have likely resulted from gene fusion events— e.g. *purH*^44^ and *histb*^45^. It is thus likely that these gene fusion events occurred after the individual gene products had been selected for higher *k_cat_*s. Gene fusions are highly polyphyletic^46-48^, a finding that supports this idea.

Further genes associated with jump behavior catalyze multiple reactions using the same binding site— e.g., *fabG* (Supplemental Table 1). Kacser and Beeby (1984)^31^ discussed the effect of such multifunctional enzymes for a scenario of highly un-specific proto-enzymes, where gene duplication becomes necessary to render increased specificity adaptive. Nevertheless, the mechanism Kacser and Beeby (1984)^31^ proposed requires assumptions about how mutations affect each catalytic activity, where experimental data indicates that such effects have to be studied on a case-by-case basis^30^. For the case of multifunctional enzymes that result from gene fusion events, independent mutation effects on both active sites seem a reasonable assumption.

### Model assumptions

A variety of sources of uncertainty make it difficult to predict experimental *k_cat_* data with the *ab initio* approach we present. Condition-dependent metabolite levels and enzyme affinities will affect enzyme saturation where our model assumes full saturation, although metabolomics data indicates that our assumption is close to reality under specific conditions^49^. Furthermore, our model has to make an assumption about the identity of biophysically constrained reactions. While EC numbers serve as a first approximation for estimating this set, there is still a high level of uncertainty in its true identity. Encouragingly, sensitivity analyses indicate that the qualitative adaptation dynamics and final growth rate are robust against the identity of the set (Supplemental Figure 5). As studies shed more light on the nature of intragenic fitness landscapes^50^, it will be valuable to model the relative contribution of intergenic and intragenic diminishing returns in more detail. Other sources of uncertainty lie in the choice of selective environments and the shape and parameters of the distribution of mutation effects. Again, sensitivity analyses show that our results are robust against these factors (Supplemental Figure 6 and Supplemental Figure 7).

### Model validation

To validate the assumptions of our modelling approach we compared model predictions to *in vitro* and *in vivo* datasets. Despite the source of uncertainty listed above and the high level of noise in the experimental data (see ^15^ for discussion) we found a significant agreement with *in vitro* data and *in vivo* estimates, where the model explained about 45% of the observed variance in *in vivo k_cat_*s. *In vitro k_cat_*s were shown to correlate with enzyme molecular weight and reaction flux (R=0.22 and R=0.45, respectively;^4^). Similarly, predicted *k_cat_*s in our model for diverse environments are correlated with enzyme molecular weight (*R*=0.28, *p*<4.4e-6) and with the mean of fluxes of parsimonious FBA ^51^ across carbon sources (*R*=0.64, *p*<2.2e-16). As our model uses information on reaction mechanisms only for the identification of the biophysically constrained set, this agreement confirms the hypothesis that these correlations are induced by differential selection pressures^4,15^. Surprisingly, we found agreement not only by correlation, but also by magnitude (Figure 4A). This finding is consistent with the realistic growth rates to which the adaptation process converges (Figure 2). The *in vivo* data used is based on quantitative proteomics data and flux estimates that assume growth maximization^37^. The better agreement of our simulations with *in vivo* data might be due to the latter being less noisy than *in vitro* estimates, but *in vivo* data could also be biased to prefer our model-based predictions. Sensitivity analyses (Supplemental Figure 5, Supplemental Figure 6) and our minimal model (Supplemental Information 2) show that the magnitude of evolved *k_cat_*s can depend on the size of the evolving set, the distribution of mutational effects, and the magnitude of biophysical constraints. We thus provide a consistent set of these parameters, but additional data is required to confirm this parameter set in the future.

## Conclusion

The prediction of evolutionary outcomes is an ultimate goal in evolutionary biology^9^. The model we present predicts data on *k_cat_* in terms of correlation and magnitude, showing that evolutionary long-term end points of *k_cat_* evolution can be predicted using systems models with unexpected accuracy, especially given the sources of model uncertainty listed above. The model predicts that diminishing returns epistasis keeps *k_cat_*s—and thus fitness—far from the global optimum, indicating the potential of engineering strategies for more efficient enzymes. While we chose *E. coli* as a model organism to study *k_cat_* evolution, the patterns we find are likely to generalize across the tree of life, where organisms with smaller effective population size than *E. coli* can be expected to show an even stronger mark of insufficient selection in their catalytic properties.

Optimality assumptions are a promising tool for understanding complex biological systems, but finite population sizes and epistatic interactions can render individual molecules far from theoretical optima— even when the underlying fitness landscape is smooth. Seeing cells through the systems perspective and modelling evolutionary history can be crucial for understanding cell behavior, as is the case for kinetic turnover numbers.

## Methods

### Growth rate predictions using MOMENT

In the simulation of kinetic parameter evolution, the growth rate that results from a given vector of turnover rates *κ* is predicted using the MOMENT algorithm^4^. MOMENT is conceptually similar to flux balance analysis (FBA^52^), in that it maximizes the growth rate *μ* by maximizing flux into a biomass reaction (*v_z_*) given a set of constraints (*v_min_* and *v_max_*):

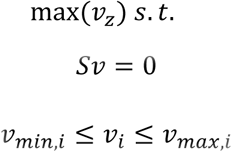

Here, *S* represents the stoichiometric matrix and *v* the vector of fluxes. MOMENT extends FBA by introducing enzyme concentrations as model variables (*g*, *mmol*/*gDW*) and recursively parsing gene-protein-reaction (GPR) rules to obtain upper limit constraints on metabolic fluxes:

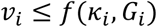

Where *G_i_* represents the set of genes involved in catalyzing reaction *i*. The respective GPR is parsed by using the maximum of enzyme concentrations to represent AND relations and the sum to model OR relations. Finally, the total weight of the metabolic proteome (*C*, *g_protein_*/*g_DW_*) and the respective enzyme molecular weights (*MW*) are used to constrain enzyme concentrations:

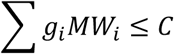

MOMENT was used to simulate growth in *i*JO1366, a genome-scale model of *E. coli* K-12 MG1655 metabolism^29^. Enzyme molecular weights were calculated based on the *E. coli* K12 MG1655 protein sequences (NCBI Reference Sequence NC_000913.3), and *C* was set to 0.32 g_protein_/g_DW_ in accordance with the *E. coli* metabolic protein fraction across diverse growth conditions^4,53^. Linear programming problems were constructed using the R^54^ packages sybil^55^ and sybilccFBA and solved using IBM CPLEX version 12.7. The growth rate *μ* (compare Figure 1) can then be obtained as the flux into the biomass reaction *v_z_*.

We classify a reaction as contributing to *in silico* growth using flux variability analysis^36^. When either the maximal flux or the absolute minimal flux through a reaction that still optimizes the growth rate *μ* in FBA is larger than 10^−6^ mmol gDW^−1^ h^−1^, we call a reaction “contributing to growth *in silico*”.

### A Markov Chain Monte Carlo algorithm for simulating *k_cat_* evolution

We assume a genetically homogenous population of cells with a population size equal to the effective population size estimated for *E. coli* (*N_e_* = 2.5e7,^33^). A single iteration of the Markov Chain Monte Carlo algorithm starts as follows: A mutation affecting the *k_cat_* of a single randomly chosen enzyme *i* is simulated as multiplying an original *k_cat_* (=*κ_i_*) by a factor α that is drawn from a lognormal distribution with mean and standard deviation in log scale log(3/2) and 0.3, respectively. This distribution determines the jump size in the space of *k_cat_*s, but not the ratio between deleterious to advantageous mutations (see below).

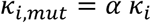

The growth rate of the original strain (*μ*) and the strain carrying the mutation affecting *κ_i_* (*μ_mut_*) is then calculated by solving the MOMENT problem detailed above (also see Figure 1). Assuming that fitness is proportional to growth rate, we can obtain the selection coefficient *s* and the fixation probability *π*^34^:

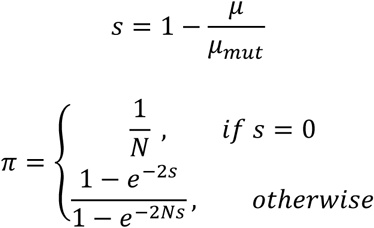

The fixation probability *π* is then used to decide the fixation of the novel mutation. In case of a successful fixation event the vector of *k_cat_*s *κ* is then updated at position *i* with the newly fixed mutation, or, in case of an unsuccessful fixation event, the previous *κ_i_* remains the most abundant allele. The next iteration of the algorithm starts with introducing a novel change in the *k_cat_* of a random enzyme, and so on. A typical simulation run simulates around 10^8^ mutations that have the chance to become fixed, requiring 10^8^ linear programs to be solved for a single replicate.

The high population size allowed us to optimize simulation performance by heuristically setting the ratio of deleterious to advantageous mutations: the growth rate for a deleterious mutation was simulated once, but their fixation was sampled multiple times to arrive at a 100:1 ratio between deleterious and advantageous mutations. Certain reaction mechanisms were shown to consistently exhibit low *k_cat_*s^15^. We use the enzyme commission (EC) number to set the reactions belonging to the three (out of six) top level codes with the highest median *in vitro k_cat_*–namely oxidoreductases, hydrolases, and isomerases–as biophysically unconstrained. In order to allow an unbiased comparison to experimental data, all reactions for which data was available were also set as unconstrained. The remaining reactions were considered biophysically constrained and were fixed to the median of *in vitro k_cat_* measurements (13.7 s^−1^). The *k_cat_*s of unconstrained reactions were initialized to 10^−3^ s^−1^.

In order to simulate diverse environments, we applied random sampling of a new environment every 1000 iterations. Here, oxygen uptake was allowed with probability ½, and the environment always contained at least one randomly chosen source of each carbon, nitrogen, sulfur, and phosphate. A number of additional sources were drawn from a binomial of size 2 with success probability 1/2. This process was repeated until a growth sustaining environment was found and the following 1000 mutations were simulated in this novel environment.

### Statistics

Pearson’s *R* was used to test for significant correlation with a two-sided t-test as implemented in the cor.test() function of the R environment^54^.

## Acknowledgements

The authors would like to thank Abdelmoneim Desouki for his support in using the sybilccFBA package, and Ron Milo and Laurence Yang for helpful discussion.

This research used resources of the National Energy Research Scientific Computing Center, a DOE Office of Science User Facility supported by the Office of Science of the U.S. Department of Energy grant number DE-SC0008701.

This work was supported by the Novo Nordisk Foundation Grant Number NNF10CC1016517.

## Contributions

DH, DCZ, and BOP designed the study. DH conducted all modelling, simulation, and data analysis. DH, DCZ, and BOP wrote the paper.

